# Structural effects driven by rare point mutations in amylin hormone, the type II diabetes-associated peptide

**DOI:** 10.1101/2021.02.17.431675

**Authors:** Wendy S. Mendes, Octavio L. Franco, Sergio A. Alencar, William F. Porto

**Affiliations:** Programa de Pós-Graduação em Ciências Genômicas e Biotecnologia, Universidade Católica de Brasília, Brasília-DF, Brazil; Centro de Análises Proteômicas e Bioquímicas, Pós-Graduação em Ciências Genômicas e Biotecnologia, Universidade Católica de Brasília, Brasília-DF, Brazil; S-Inova Biotech, Pós-Graduação em Biotecnologia, Universidade Católica Dom Bosco, Campo Grande, MS, Brazil; Porto Reports, Brasília-DF, Brazil http://www.portoreports.com

**Keywords:** Molecular dynamics, Islet amyloid, S20G mutation, Pramlintide

## Abstract

**Background:** Amylin is a 37-amino-acid peptide hormone co-secreted with insulin, which participates in glucose homeostasis. This hormone is able to aggregate in a β-sheet conformation and deposit in islet amyloids, a hallmark in type II diabetes. Since amylin is a gene-encoded hormone, this peptide has variants caused by point mutations that can impact its functions.

**Methods:** Here, we analyzed the structural effects caused by S20G and G33R point mutations which, according to the 1000 Genomes Project, have frequency in East Asian and European populations, respectively. The analyses were performed by means of aggrescan server, SNP functional effect predictors, and molecular dynamics.

**Results:** We found that both mutations have aggregation potential and cause changes in the monomeric forms when compared with wild-type amylin. Furthermore, comparative analyses with pramlintide, an amylin drug analogue, allowed us to infer that second α-helix maintenance may be related to the aggregation potential.

**Conclusions:** The S20G mutation has been described as pathologically related, which is in agreement with our findings. In addition, our data suggest that the G33R mutation might have a deleterious effect. The data presented here also provide new therapy opportunities, whether for creating more effective drugs for diabetes or implementing specific treatment for patients with these mutations.

**General Significance:** Our data could help to better understand the impact of mutations on the wild-type amylin sequence, as a starting point for the evaluation and characterization of other variations. Moreover, these findings could improve the health of patients with type II diabetes.

## 1 Introduction

Amylin is a hormone co-secreted with insulin at a rate of 1:100 by the pancreatic β-cells [1]–[3]. It participates in glucose homeostasis, causing a number of effects, including reduction in food intake, nutrient delivery delay, muscle glycogen synthesis inhibition and lipolytic-like effects [1]. Encoded from the *IAPP* gene, amylin has a precursor and a pro-form, with the precursor being synthetized with 89 amino acid residues [3]. This precursor loses the signal peptide, resulting in the 67-residue pro-form, which is processed to generate the 37-amino-acid amylin mature form [4] (Fig. 1A). This peptide is part of the calcitonin gene peptide superfamily [5] and is encoded, in humans, on chromosome 12p12.1 [3].

**Fig. 1.**
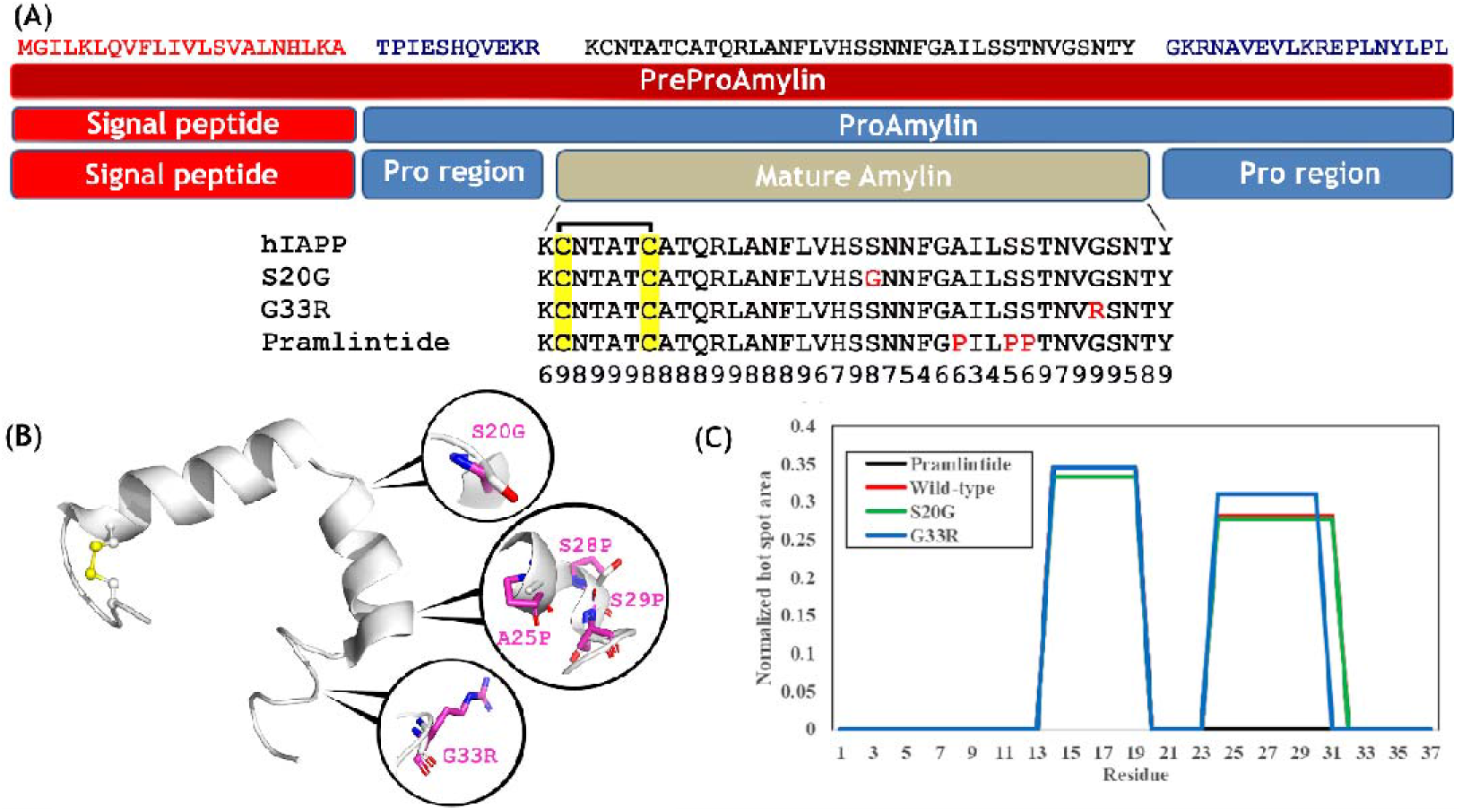
Amylin and its variants. (A) Amylin processing and alignment with its variants. PreProAmylin is the amylin precursor with 89 amino acids. The precursor has a 22-residue signal peptide, which is cleaved to form ProAmylin. This propeptide has two flanking regions that are removed by the prohormone convertases enzymes PC2 and PC3 in the process to release the mature peptide. In the process, the peptidyl amidating mono-oxygenase complex acts by catalyzing an amidation reaction on the C-Terminal. The 37 amino acid residues mature amylin has a disulfide bond, formed by Cys^2^ and Cys^7^ (highlighted in yellow). The SNPs S20G and G33R are the only missense variants reported with frequency in the 1,000 genomes project. The amylin analogue drug, pramlintide, has the point mutations A25P, S28P and S29P derived from rat amylin, to reduce the molecule aggregation capacity. The amino acids that differ from the wild-type human amylin in the alignment are highlighted in red. The numbers below the alignment represent an increasing scale of conservation from Consurf (see Fig S1). (B) Tridimensional structure of wild-type amylin. This molecule structure (PDB 2l86) present two α-helices in Thr^6^-His^18^ and Ser^20^-Ser^29^. The disulfide bond is represented in ball and stick. The approximate images show the amino acid variants in positions 20, 25, 28, 29 and 33, showing the mutation residue side chains in S20G, pramlintide, and G33R (magenta) superimposed with the original side chains (white). The results from the validation of models are available in Table S2. (C) Normalized hot spot area (NHSA) graphic obtained using aggrescan software. The NHSA is calculated by the hot spot area divided by the number of residues in the sequence. The hot spot is determined by a region with residues that passes a calculated limit and respects the ‘no proline’ condition established, because this amino acid is an aggregation breaker. These data indicate the wild-type has two distinct regions of aggregation in 13-20 and 23-32 residues. The drug pramlintide, well known to be less amyloidogenic and with three prolines located in the second region, has just a 13-20 residue region. Although the variants also have two regions with high hotspot area values, the second region from G33R only includes residues 23-31.

This hormone is often related to the development or aggravation of type II diabetes [6]. Amylin was first isolated from the islet amyloid in patients with non-insulin-dependent diabetes mellitus [3], and, therefore, amylin is also known as the islet amyloid polypeptide. The islet amyloid is a denomination for the deposition of amylin fibrils, an aggregate polypeptide state with several β-sheet conformations [1], [7]. This amyloid form is related to cytotoxicity. However, its mechanisms are still not fully understood [8]. Even though the islet amyloid is a hallmark in type II diabetes, there is a hypothesis that the oligomeric states could be the toxic form in amylin, and the process of deposition is an immune response to prevent damage [1], [9].

Comparisons involving amylin from different species and even with other peptides from the calcitonin superfamily have enabled a better understanding of some regions in this molecule [2], [10]. Analyzing the conservation between these sequences and the aggregation response, the region harboring the amino acids Ser^20^-Ser^29^ was suggested as the main one responsible for the aggregation function [2], [8]. However, mutations within and outside this region are capable of affecting amyloid properties [8]. The only substitution described with amyloid effect *in vivo* is the S20G in the amylin mature peptide [11]. Since the existence and maintenance of the amidated C-Terminal and the Cys^2^-Cys^7^ disulphide bond are essential to amylin bioactivity [12], the hormone’s activity is also susceptible to other mutations.

The study of mismatches in homologous sequences enables the development of synthetic peptides with specific properties. In this context, the drug pramlintide was developed based on a non-amyloidogenic rat amylin sequence, which diverges in 7 residues from the human mature amylin sequence [10]. This amylin analogue drug has three mutations (A25P, S28P and S29P) and aggregates less than human amylin. Pramlintide also mimics some of the amylin functions, such as gastric emptying delay, and it is used as pharmacological treatment for some type II diabetic patients [13].

Nowadays, computational tools have been used for the evaluation of variant effects [14], with a focus on mutations with frequency in the population, described in projects such as the 1,000 Genomes Project [15]. The prediction tools are able to predict some deleterious behavior by means of different strategies that can be classified in four groups, which are sequence homology-based, supervised learning-based, structure-based and consensus-based [16]. Although these tools have limitations described in benchmarking analyses [17]–[20], they still can be used as a filter or indicator in studies on the effect of single-nucleotide polymorphisms (SNPs) [16]. Then, to improve the accuracy of predictions, other techniques, such as molecular dynamics, can be used [21]–[25]. This approach has been used by our group to study other mutations related to peptide hormones, such as guanylin [26], uroguanylin [27], lymphoguanylin [28] and growth hormone receptor [29]. Our hypothesis is that mutations in the amylin gene can alter the peptide structure. Therefore, we aim to analyze these structural variants. Here we show that rare mutations in mature amylin caused by SNPs are located in conserved positions, maintaining the aggregation potential and a similar structure conformation to the wild-type after an unfolding dynamics simulation. Despite that, we also observed that variants cause changes in the monomeric forms when compared to the wild-type.

## 2 Results

### 2.1 Amylin mutations from European or Asian populations are in highly conserved positions

The amylin hormone is coded by its gene as a precursor protein and undergoes several processing steps until the release of the 37-amino-acid mature peptide (Fig. 1A); therefore, this work focused on mutations in the mature chain. According to the 1000 Genomes data, the S20G mutation had frequency in the East Asian population (0.011) and was reported in three Chinese and one Japanese subpopulation, while the G33R mutation had frequency in the European population (0.001), reported in a British subpopulation. The positions where the mutations occur were classified as conserved according to Consurf’s amino acids evolutionary conservation evaluation (Fig. 1A and Fig. S1). According to the pairwise F_ST_, both mutations presented no genetic differentiation in their respective populations (Table S1).

### 2.2 European rare mutation is predicted as deleterious

The G33R mutation was predicted as deleterious in all five supervised learning method-based tools, and in four out of five sequence-based tools, while only one tool of the consensus-based group had a deleterious prediction (Table 1). In contrast to the S20G mutation with only three deleterious prediction results, two of them were in tools that are part of the sequence-based group, and the other in a tool based on a supervised learning method (Table 1). The consensus-based prediction for the S20G mutation resulted in none of these tools with a deleterious prediction (Table 1). As well as the structure-based tools that evaluate the stabilizing effect, three out of five predicted G33R as a stabilizing mutation, while only one out of five had the same prediction for S20G (Table 1).

**Table 1.** Prediction results of amylin SNPs analyzed by 20 prediction tools classified in four different groups

| SNP rs # | Amino acid change<sup>a</sup> | Sequence-based<sup>b</sup> |  |  |  |  | SLM-based<sup>b</sup> |  |  |  |  | Consensus-based<sup>b</sup> |  |  |  |  | Structure-based<sup>c</sup> |  |  |  |  |
| --- | --- | --- | --- | --- | --- | --- | --- | --- | --- | --- | --- | --- | --- | --- | --- | --- | --- | --- | --- | --- | --- |
|  |  | SIFT | PROVEAN | Mutation Assessor | Panther | FATHMM | SNAP | SuSpect | MutPred | PolyPhen-2 | PhD-SNP | Condel | Meta-SNP | SNPeffect | PredictSNP | TransFIC | SDM | PoPMuSiC | HOTMuSiC | SNPMusic | Fold-X |
| rs1800203: A>G | S20G | N | D | N | D | N | D | N | N | N | N | N | N | N | N | N | 0.16* | 0.41 | -0.75 | -0.83 | 0.12 |
| rs200376097: G>A | G33R | D | D | D | D | N | D | D | D | D | D | N | N | N | D | N | 0.35* | -0.17* | 1.27* | 0.43 | 2.15 |

<sup>a</sup> Both SNPs were validated by frequency in 1,000 Genome Project.
<sup>b</sup> N: Neutral; D: Deleterious.
<sup>c</sup> These prediction tools present the results in the form of a score (represented in the table) that can be interpreted as stabilizing or destabilizing. The results interpreted as stabilizing are marked with an asterisk (\*) after the number. The interpretation of whether the stabilizing effect is deleterious or neutral is further discussed.
In the analyses the mature amylin peptide structure was used (PDB ID: 2L86).

### 2.3 Aggregation and secondary structure are maintained in the variants

Aggrescan analysis showed two aggregation regions in the wild-type sequence, which were maintained in both mutations. However, pramlintide, known to aggregate less than human amylin, loses the second region (Fig. 1C).

The resolved native amylin structure is composed of two α-helices (Fig. 1B). Structurally, the selected mutations, S20G and G33R, are located in the second α-helix and in the C-Terminal loop, respectively (Fig. 1B). The validated models (Table S2) for the variants and for pramlintide used the native amylin structure as a template.

The molecular dynamics simulations were performed in saline solution, while the NMR structure used as a template (PDB ID: 2L86) was resolved in SDS micelles [30]. Hence, we can observe the unfolding behavior in molecular dynamics, with the backbone’s RMSD reaching values higher than 4 Å, indicating several conformation changes in all models (Fig. 2A). Regarding the variants, the G33R mutation reached the highest radius of gyration (Rg) and solvent accessible surface area (SASA) values between all those in the ensemble in the first half of the simulation period (Fig. 2B and C). Although the S20G mutation kept a similar structure to wild-type, its Rg and SASA values were similar to those of pramlintide (Fig. 2B and C). RMSF variations between the wild-type and the variants were notable, especially in the G33R, with higher values in at least 3 regions of the sequence (Thr^4^-Gln^10^, Val^17^-Asn^22^, Asn^31^-Tyr^37^), including the mutant residue position (Fig. 2D). The S20G variant maintained a similar fluctuation to wild-type in the mutant residue position (Fig. 2D).

**Fig. 2.**
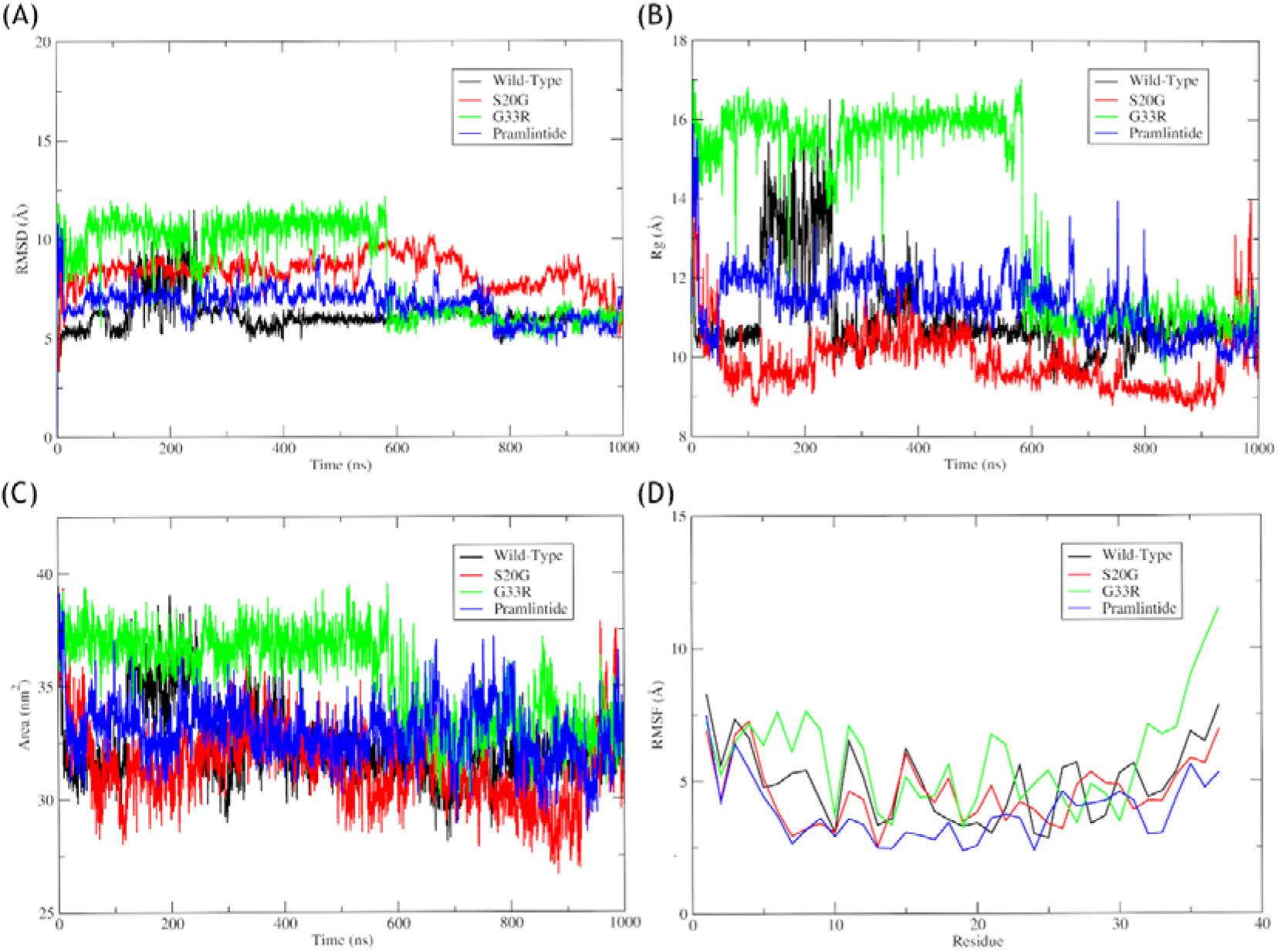
Analyses of molecular dynamics trajectories. (A) The RMSD variation during the simulations. Due to values higher than 4Å, we notice a structure change in all models. (B) The radius of gyration (Rg) variation during the simulations. (C) The solvent accessible surface area (SASA) variation during the simulations. The Rg and the SASA have similar behavior, showing the structure forms with higher expansion also have high solvent accessible surface area. (D) The RMSF variation between the ensembles. Gly^20^ introduction in the S20G has not notably altered RMSF in comparison with the wild-type values. The movement in the final loop (T^30^-Y^37^) in G33R values increased, probably due to Arg^33^ introduction. In the same region, the drug pramlintide has lower values than wild-type, a possible reflection of Pro^25^, Pro^28^ and Pro^29^ addition.

In the principal component analysis (Fig. 3), two very distinct clusters were observed in the G33R mutation, where a long α-helix in the distinct cluster was noted in the same time period as the Rg and SASA peak (Fig. 2B and C). Despite the changes suffered after 1 μs in an unfolding dynamics simulation, the variants maintained a similar structure to the wild-type (Fig. 3). Although the wild-type and the variant structures are similar, they have differences in the length of their α-helices and the variants’ eigenvectors projection clusters were mainly overlapped with the pramlintide structure (Fig. 3). Structurally, pramlintide was capable of maintaining only the N-Terminal α-helix (Fig. 3).

**Fig. 3.**
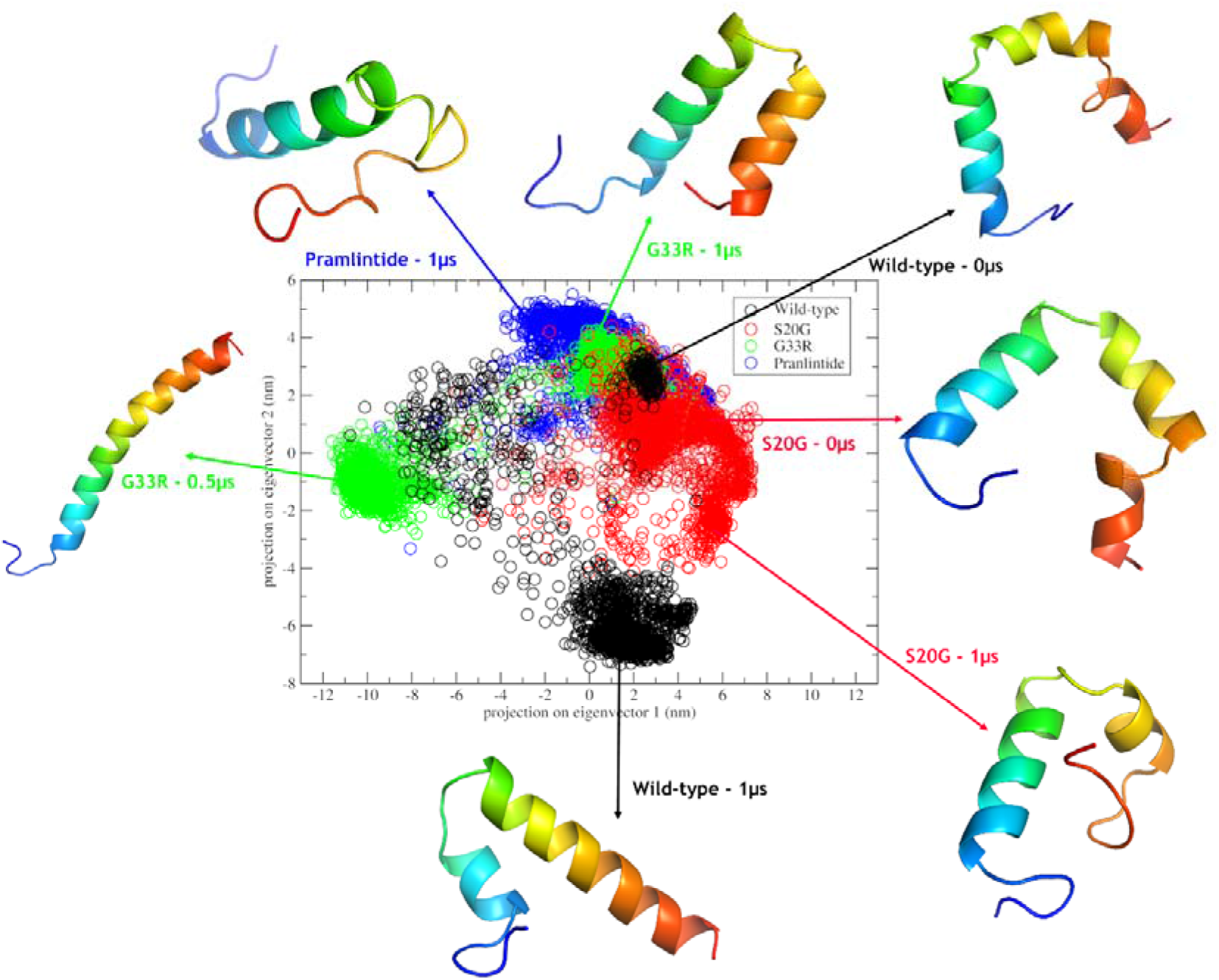
Eigenvector clusters’ representative structures. The four models were submitted to an unfolding dynamics simulation (See Material and Methods) and had its principal component analyzed. The representative structures from clusters formed by projection of the motion of amylin in phase space along the first two principal eigenvectors are presented. The arrows have a color corresponding to the graphic. The structures are colored in a rainbow pattern with the N-Terminal in blue and the C-Terminal in red. The variants and the drug pramlintide have a small overlap among the structures, while most wild-type structures form a distant cluster (eigenvector 1: 2 and eigenvector 2: −6). The G33R left cluster (eigenvector 1: −10 and eigenvector 2: 1) represents an intermediate structure from the 0 – 600 ns dynamics period.

## 3 Discussion

Amylin is a peptide hormone that participates in glucose homeostasis and, due to the amyloidosis phenomenon, has a key role in type II diabetes pathophysiology [31], [32]. In fact, the amyloidogenic characteristic has been extensively discussed [1], [7], [33], [34], and is understood as essential to determine normal or pathological peptide behavior [32].

Because amylin is a gene-encoded hormone, its amino acid sequence can be modified by SNPs on the *IAPP* gene. Many strategies for SNP discovery have been described, but none of them is flawless [35]. Using SNPs with reported frequency means the variants were found in the population and they were already able to affect people. Thus, the understanding of the impact that mutations with frequency cause is an immediate need, making it necessary to create or adapt therapeutic intervention to avoid and remediate the possible issues in these affected populations.

The rare variants selected here were caused by SNPs with global frequencies lower than 1%. The evolutionary maintenance of the amino acid signifies that the amino acid is probably needed in function or structure in the peptide, and the S20G and G33R mutations were found in positions classified as evolutionarily conserved (Fig. S1). As matter of fact, amino acid residues that perform specific functions, such as catalytic functions and interactions, are often more conserved [36], [37]. Although this suggests conserved mutations have a higher propensity to exert functional or structural effects, the relationship between pathogenicity and conservation must be carefully analyzed [38].

To contribute to the determination of pathogenicity we analyzed the prediction of SNP effect, but this, specifically, must be carefully analyzed because benchmarking analyses have shown these predictors lose accuracy to specific proteins [17]–[20]. Due to the known prediction limitations, other analyses were performed, such as aggregation prediction and molecular dynamics. The G33R mutation has a highly deleterious prediction (Table 1) and, until now, this mutation has no effects reported in the literature. However, the S20G mutation is mostly neutral-predicted (Table 1), although its deleterious effect has been shown [11], [39], [40], and, therefore, the predictions for the S20G mutation could be false negatives.

Other interesting analysis concerns the prediction tools that evaluate effects on molecule stability (Table 1). We can observe that G33R was mainly predicted as a stabilizer, while the S20G results were often not stabilizing. The amyloidosis process is dynamic, reversible and directed to fibril formation; therefore, more stable molecules usually hinder a reversal of the process with the oligomeric formation that is believed to be amylin’s toxic form [9].

The S20G mutation is described as involved in type II diabetes and has increased aggregation capacity [11], [39]–[41], which was demonstrated to be due to structural changes and the existence of three subunits in a fibril, compared to wild-type fibrils with only two subunits [7]. The accommodation of three subunits in a fibril could be related to the mutant amino acid being a glycine, an achiral amino acid. This suggestion is supported by Rg and SASA data, in which S20G values are slightly lower than the wild-type ones, representing a comparatively more compressed conformation state. Interestingly, Asians have higher type II diabetes rates [42], which might be related to the S20G frequency in the East Asian population. On the other hand, the G33R mutation has a unique structural form (Fig. 3) caused by a repulsion between the Arg^11^ and the C-Terminal portion, which became cationic due to Arg^33^ inclusion. The formation of this long α-helix coincides with the peak of the G33R Rg and SASA values and exposes a molecule with broader movement for this structure, caused by the addition of arginine. Furthermore, the substitution of a glycine, which is apolar and achiral, by an arginine, with its positive charged side chain, could impact the G33R fibril formation. This α-helix structural form could also affect the peptide’s function.

Point mutations are able to affect the aggregation capacity, *e.g.,* the drug pramlintide has three variants, A25P, S28P and S29P, near the C-Terminal, and reduced amyloidogenicity. The difference between pramlintide and wild-type amyloidogenicity, along with the fact that amyloidosis capability is important to amylin evaluation, motivated us to use the pramlintide drug sequence as a control in the analyses with the wild-type sequence and the mutations (Fig. 1A). Our data show that for both S20G and G33R mutations, the aggregation potential is maintained (Fig. 1C), and these mutations also cause structural behavior changes in the monomeric forms when compared with the wild-type (Figs. 2 and 3). These data support what has been described for the S20G mutation, and also suggest a pathological potential for the G33R mutation. Thus, the aggregation potential is maintained in G33R (Fig. 1C) and so this variant could be involved in type II diabetes pathophysiology, increasing the formation of amyloid deposits.

Interestingly, the second region of aggregation and the C-Terminal α-helix present in both variants and in the wild-type are absent in the pramlintide sequence. This observation is related to the replacement of three amino acids by proline residues in positions 25, 28 and 29 of the drug pramlintide, since proline residues are known to be α-helix breakers. Therefore, we can infer that the existence of this secondary structure is probably related to the aggregation capacity. There could be a relationship between aggregation potential and α-helix maintenance (Fig. 1C and Fig. 3), which could act as a classifier for amyloidogenic potential, with the possibility of being used as an indicator of amylin’s deleterious mutation.

The existence of SNPs in a sequence can be related to composition and structural effects [43]. Non-synonymous mutations, for instance, represent an amino acid change with several potential effects on structure, interactions, and conformations, even though with different disease severity [44]. That way, a structural study can be a valuable tool in analyzing the effects the mutations may have on protein function or physiology, a strategy widely used in previous studies with other proteins [16], [21], [22], [24]–[26]. This type of study is usually limited by the need for a resolved structure. However, due to recent advances in the field of computational three-dimensional structure prediction, this limitation might be overcome. The program AlphaFold2, an accurate method of structure prediction based on a deep-learning method, has improved structural predictions, with some results indistinguishable from structures determined by experimental methods [45]. Nevertheless, the AlphaFold2 limitations are still not clear, including cases of very dynamic structures, such as amylin, which change upon oligomerization (recently, cryo-em experiments demonstrated changes in the oligomeric conformation caused by the S20G mutation [7]). Even with a possible limitation, the AlphaFold2 breakthrough could represent an immeasurable improvement in future studies.

## 4 Conclusions

The mature human amylin sequence has only two missense mutations with reported frequency in the 1,000 Genomes Project, the S20G and the G33R mutations. There are some studies that try to describe amylin cytotoxicity, and others that try to evaluate whether these mutations influence this cytotoxicity or even whether they play a pathological role in other forms. Structural studies can combine both fields and try to identify the relationship between the variation and the deleterious effect, as well providing a better understanding of peptide behavior.

In this context, structural analysis demonstrated here that both mutations probably cause changes in the monomeric forms in comparison to the wild-type. Furthermore, a unique structure form in the G33R mutation was also observed, and its biological effects are still unknown. Data presented here also show aggregation potential was maintained in the structures with S20G and G33R mutations, and that this capacity is probably related to the maintenance of the C-Terminal α-helix. This work allows a better understanding of the structural effects caused by *IAPP* SNPs and could improve future SNP evaluation. Therefore, better comprehension of how mutations influence the wild-type amylin sequence may allow new therapeutic opportunities, including creating more effective drugs for type II diabetes or implementing specific treatment for the patients with these mutations. Our findings can also help in continuing the evaluation and characterization of other variations.

## 5 Material and Methods

### 5.1 Databases and SNP selection

The SNPs were searched in the dbSNP database (build 153) using the Variation Viewer navigator from the NCBI (https://www.ncbi.nlm.nih.gov/variation/view/) [46]. The missense variant filter was applied and only the SNPs located in the mature sequence with frequency in the 1,000 Genomes Project phase 3 (https://www.internationalgenome.org/), reference assembly GRCh38 [15] were selected. The amylin sequence and the protein structure files of the mature peptide (PDB ID: 2L86) were obtained from the RCSB Protein Data Bank [30], [47].

### 5.2 SNP frequency

The SNP frequency data of the *IAPP* gene were obtained from the 1,000 Genomes Project (phase 3) browser [15]. The browser provides the frequencies of all SNPs identified in the genomes of 2,504 individuals from 26 populations obtained through a combination of low-coverage (2–6×) whole genome sequence data, targeted deep (50–100×) exome sequencing, and dense SNP genotype data. The 26 populations studied were grouped by the predominant component of ancestry into five super-populations: African (AFR; 661 individuals), Admixed American (AMR; 347 individuals), East Asian (EAS; 504 individuals), South Asian (SAS; 489 individuals), and European (EUR; 503 individuals). The fixation index (F_ST_) was measured according to the model developed by Wright [48].

### 5.3 Functional analyses of variants

The SNPs were submitted to functional impact prediction using 20 computational tools, divided into 4 different groups, as described below.

#### 5.3.1 Sequence homology-based methods

These methods determine whether a variation has functional effect or not by means of homologous sequence multiple alignments and sequence conservation. Sorting Intolerant From Tolerant (SIFT) [49], Protein Variation Effect Analyzer (PROVEAN) [50], Mutation Assessor [51], Protein Analysis Through Evolutionary Relationships (PANTHER) [52] and Functional Analysis through Hidden Markov Models (FATHMM) [53] predictors were applied. We used the SIFT sequence option in the single protein tools; and the PROVEAN protein tool in the PROVEAN web server; the remaining tools were used in default settings.

#### 5.3.2 Supervised learning methods (SLM)

Tools such as Screening for Non-Acceptable Polymorphisms (SNAP) [54], Disease-Susceptibility-based SAV Phenotype Prediction (SuSPect) [55], MutPred [56], Polymorphism Phenotyping v2 (PolyPhen-2) [57] and Predictor of human Deleterious Single Nucleotide Polymorphisms (PhD-SNP) [58] use an algorithm to learn patterns of conservation and/or physicochemical properties. In SuSpect, we analyzed the position of mutation residues in the precursor sequence using Mutate an entire protein option; the other tools were used with the default settings.

#### 5.3.3 Structure-based methods

This group analyzed the SNPs effects by means of structural information, for instance solvent accessibility and free energy variation. We used the Site Directed Mutator (SDM) [59], Fold-X [60], PopMuSiC [61], HOTMuSiC [62] and SNPMusic [63] predictors in this category. Default settings were used in the first two tools and in the last three, the manual approach was used to select the specific SNPs. In SDM, the negative and positive values represent a destabilizing and stabilizing prediction, respectively [59]. In FoldX, positive results are usually indicative of problems with the structure [60]. In PoPMuSiC and HoTMuSiC, negative and positive values, respectively, mean a stabilizing mutation. Finally, in SNPMuSiC, positive and negative scores correspond to a deleterious and neutral prediction.

#### 5.3.4 Consensus-based methods

The predictors in this category combine methods and tools to create a SNP effect classifier. In this group, the following tools were used: Consensus Deleteriousness (Condel) [64], Meta-SNP [65], SNPeffect [66], Transformed Functional Impact for Cancer (TransFIC) [67], [68] and PredictSNP [69]. The last one used the precursor FASTA sequence as an input, and the tools nsSNPAnalyzer and PANTHER were included in the consensus. Default settings were used in the remaining predictors.

### 5.4 Evolutionary conservation analysis

The Conservation Surface-mapping (ConSurf) server is a tool that, based on the phylogenetic relations between homologous sequences, can estimate the evolutionary conservation of amino acid positions in a protein molecule [70]. Using the precursor sequence in FASTA format obtained from the National Center of Biotechnology Information (NP_000406.1), ConSurf, in ConSeq mode, carried out a search for close homologous sequences using CSI-BLAST (3 iterations and 0.0001 e-value cutoff) against the UNIREF-90 protein database [71], [72]. The parameters were set as follows: maximum number of homologs as 150 and a 35 minimal to 95 maximal percentage ID between sequences. Multiple sequence alignment and calculation methods were used as default (MAFFT-L-INS-I and Bayesian). The sequences were then clustered and highly similar sequences removed using CD-HIT [73]. The empirical Bayesian algorithm was used to compute the position-specific conservation scores [74].

### 5.5 Aggregation hot spot area prediction

Aggrescan is based on an aggregation-propensity scale for natural amino acids and is able to predict aggregation-prone segments in protein sequences [75]. The sequence in FASTA format was used as input; for each, aggrescan returned the aggregation-propensity value for residue and aggregation hot spots from the peptide aggregation profile.

### 5.6 Molecular modelling

One hundred molecular models for each ensemble (the two variants and the drug pramlintide) were constructed by comparative molecular modelling by means of MODELLER 9.16 [76] using the structure of amylin (PDB ID 2l86) as a template. Three-dimensional models were constructed using automodel and environ classes from MODELLER with default settings. The final models were selected according to the discrete optimized protein energy score (DOPE score), which indicates the most probable structures [77]. The best models were evaluated through PROSA II [78], QMEAN [79] and PROCHECK [80] softwares. PROSA II is a tool applied to evaluating protein structures by means of a score based on a comparison with all known protein structures that indicate the fold quality. QMEAN uses a scoring function that combines six structural descriptors to estimate the absolute quality of a protein structure. PROCHECK checks the stereochemical quality of a protein structure by means of Ramachandran plot, in which models with good quality have more than 90% of residues in the most favored and additional allowed regions. The model structures were visualized in PyMOL (http://www.pymol.org).

### 5.7 Molecular dynamics simulation

The molecular dynamics simulation of the wild-type, the variants and the drug pramlintide were performed in the GROMACS 4.6 [81] package using the CHARMM force field [82], [83]. The simulations were done in a saline environment (0.2 M NaCl) where the structures were immersed in water cubic boxes with a 1.2 nm distance between the protein and the edge of the box. Geometry of water molecules was constrained by using the SETTLE algorithm [84]. All atom bond lengths were linked by using the LINCS algorithm [85]. Electrostatic corrections were made by the Particle Mesh Ewald algorithm [86], with a threshold of 1.4 nm to minimize the computational time. The same cut-off radius was also used for van der Waals interactions. The list of neighbors of each atom was updated every 500 ps of simulation. The steepest descent algorithm was applied to minimize system energy for 50,000 steps. After that, the system temperature was normalized to 310 K for 100 ps, using the velocity rescaling thermostat (NVT ensemble). The system pressure was subsequently normalized to 1 bar for 100 ps, using the Parrinello-Rahman barostat (NPT ensemble). The system with temperature normalized to 310 K and pressure normalized to 1 bar was simulated for 1 μs by using the leap-frog algorithm as integrator.

### 5.8 Analyses of molecular dynamics trajectories

Molecular dynamics simulations were analyzed by means of the backbone root mean square deviation (RMSD), radius of gyration (Rg), solvent accessible surface area (SASA) and root mean square fluctuation (RMSF) using the g_rms, g_gyrate, g_sas and g_rmsf built-in functions of the GROMACS package. Except for the RMSD, which used GROMACS group 4 (backbone) for analyses, the functions used GROMACS group 1 (protein). Principal component analysis (PCA) was used to analyze and visualize the protein motion in the simulation. For that, the protein movement is extracted and split into different components. For visualization, the two largest classes of motion (the 1^st^ and the 2^nd^ eigenvectors) were plotted against each other. The PCA, or essential dynamics analysis, was done using the g_covar and g_anaeig functions.

## 6 Acknowledgements

This work was supported by CNPq (Conselho Nacional de Desenvolvimento Científico e Tecnológico); CAPES (Coordenação de Aperfeiçoamento de Pessoal de Nível Superior), FAPDF (Fundação de Amparo a Pesquisa do Distrito Federal) and FUNDECT (Fundação de Apoio ao Desenvolvimento do Ensino, Ciência e Tecnologia do Estado de Mato Grosso do Sul).

