## Supplementary Material for "Structural effects driven by rare point mutations in amylin hormone, the type II diabetes-associated peptide"

### 1 Supplementary information

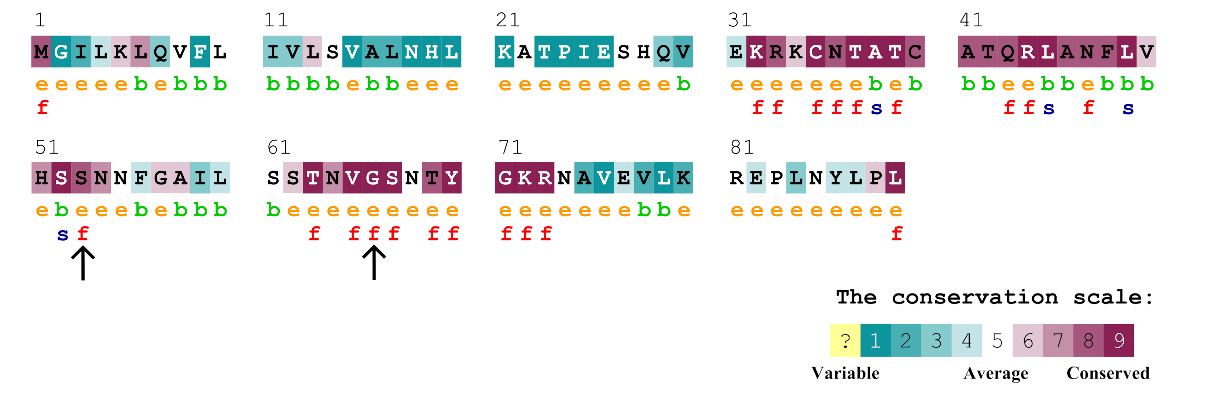

**Fig. S1. Consurf amino acid conservation pattern.** Consurf server searches for homologous sequences to the query sequence and build patterns based on amino acid conservation. The patterns are shown by a color scale, in which the grade of 1 (turquoise) represent the residues with high variability, and the grade of 9 (magenta) represent the residues that are highly conserved among homologous. The variability of an amino acid is usually associated with fundamental structural and/or functional activity. These characteristics were predicted and represented by the letters f- a predicted functional residue (highly conserved and exposed) and s- a predicted structural residue (highly conserved and buried). It was also used the letters e and b to represent residues exposed or buried, respectively, according to the neural network algorithm. In the figure, the black arrows indicate the changed residue on S20G and G33R located in the mature peptide. These SNPs have a highly conserved pattern, are exposed, and are predicted as functional residues. Hence, is expected the variations may have effects on amylin peptide.

**Table S1.** Wright's Fixation indices (F_ST_) obtained from pairwise population comparisons

| **Mutation** | **AMR-AFR** | **AMR-EUR** | **AMR-EAS** | **AMR-SAS** | **AFR-EUR** | **AFR-EAS** | **AFR-SAS** | **EUR-EAS** | **EUR-SAS** | **EAS-SAS** |
| --- | --- | --- | --- | --- | --- | --- | --- | --- | --- | --- |
| S20G | - | - | 0.005 | - | - | 0.006 | - | 0.006 | - | 0.005 |
| G33R | - | <0.001 | - | - | 0.001 | - | - | 0.001 | <0.001 | - |

**Table S2.** Results of the best models validation in the molecular modeling

| **Structure** | **Z-score**  **(Prosa II)** | **QMEAN4** | **Ramachandran plot** | |
| --- | --- | --- | --- | --- |
|  |  |  | **Most favored regions** | **Additional allowed regions** |
| S20G | -0.52 | -0.62 | 84.4% | 9.4% |
| G33R | -0.14 | -1.15 | 88.2% | 8.8% |
| Pramlintide | -0.24 | -0.81 | 90.0% | 6.7% |
